# Medullary vein architecture modulates the white matter BOLD cerebrovascular reactivity signal response to CO_2_: observations from high-resolution T2^*^ weighted imaging at 7T

**DOI:** 10.1101/2021.09.03.458842

**Authors:** Alex A. Bhogal

## Abstract

Brain stress testing using blood oxygenation level-dependent (BOLD) MRI to evaluate changes in cerebrovascular reactivity (CVR) is of growing interest for evaluating white matter integrity. However, even under healthy conditions, the white matter BOLD-CVR response differs notably from that observed in the gray matter. In addition to actual arterial vascular control, the venous draining topology may influence the WM-CVR response leading to signal delays and dispersions. These types of alterations in hemodynamic parameters are sometimes linked with pathology, but may also arise from differences in normal venous architecture. In this work, high-resolution T2^*^weighted anatomical images combined with BOLD imaging during a hypercapnic breathing protocol were acquired using a 7 tesla MRI system. Hemodynamic parameters including base CVR, hemodynamic lag, lag-corrected CVR, response onset and signal dispersion, and finally ΔCVR (corrected CVR minus base CVR) were calculated in 8 subjects. Parameter maps were spatially normalized and correlated against an MNI-registered white matter medullary vein atlas. Moderate correlations (Pearson’s rho) were observed between medullary vessel frequency (MVF) and ΔCVR (0.52; 0.58 for total WM), MVF and hemodynamic lag (0.42; 0.54 for total WM), MVF and signal dispersion (0.44; 0.53 for total WM), and finally MVF and signal onset (0.43; 0.52 for total WM). Results indicate that, when assessed in the context of the WM venous architecture, changes in the response shape may only be partially reflective of the actual vascular reactivity response occurring further upstream by control vessels. This finding may have implications when attributing diseases mechanisms and/or progression to presumed impaired WM BOLD-CVR.

## INTRODUCTION

Vascular reactivity mapping using blood oxygenation level-dependent (BOLD) magnetic resonance imaging (MRI) in combination with a vasoactive stimulus is of increasing interest for clinical investigation of a wide range of cerebrovascular diseases. Of particular interest is the relationship between the BOLD cerebrovascular reactivity (-CVR) response^1^ with white matter (WM) integrity as a means through which to better understand or predict the development of WM lesions^2-4^. A recurring assumption is that factors governing the signal characteristics in WM are arterial in origin and should be compared against the response of the gray matter (GM). It follows then that deviations from the GM reference response may indicate impairment, flow redistribution^5^, variable CO_2_ sensitivity^6^, or differences in the rate of the vascular response^7^. This work proposes that an additional component, related to how venous blood is drained from WM tissue, modulates the WM BOLD-CVR to stimulus separate from factors related to direct arterial control.

GM cortical arteries are organized centripetally (towards the center); arteries penetrate from the pial surface towards the center of the brain while branching within the cortex. In some cases, these vessels may also penetrate the sub-cortical WM. In this superficial brain region, drainage occurs centrifugally (away from the center) as associated veins carry blood back to the pial surface and eventually the superior sagittal sinus via the intra-cortical, subcortical, and superficial medullary veins^8, 9^. In deeper WM regions, blood is drained centripetally. This pattern is clear when observing the medullary veins that originate 1-2cm below the cortex and drain via the subependymal veins lateral to the ventricles (see figure 1)^10^. This organization, which distinctly demarcates superficial and deep cerebral drainage networks^8^, is sometimes connected by short anastomotic medullary veins and trans cerebral veins that link the pial and subependymal veins^8^. Collateral drainage is between these networks is limited. Moving inwards towards the ventricles, specific vein structures of the deep drainage system are classified by 4 convergence zones^11^ (see figure 1 in Taoka et. al 2017^10^ and for an overview of venous drainage in the cerebrum see the introduction of Khalatbari et al. 2021^9^). Based on high-resolution T2^*^ weighted imaging, it is possible to distinguish structural elements related to zones 2 through 4 (figure 1)^8^. The relationship between auto-regulatory control vessels and the venous structures that drain from associated tissues differs between the superficial and deep networks. The cortical and sub-cortical arterioles/capillaries run near their superficial draining veins/veinules^12^. This tight coupling is essential due to the high energy, and auto-regulatory demand of the neuronal tissue and also means that deoxyhemoglobin (dHb) mediated BOLD contrast changes in the cortex provide a relatively specific representation of local auto-regulatory control. It is also worth noting that cortical physiology leads to a several-fold increase in vascular density in superficial regions as compared to the deeper brain regions^13,14^. An additional consequence is that medullary veins draining the deep WM may pool blood from distal control vessels. In some cases, venous blood will traverse several zones of convergence before finally congregating at the subependymal veins.

**Figure 1A:**
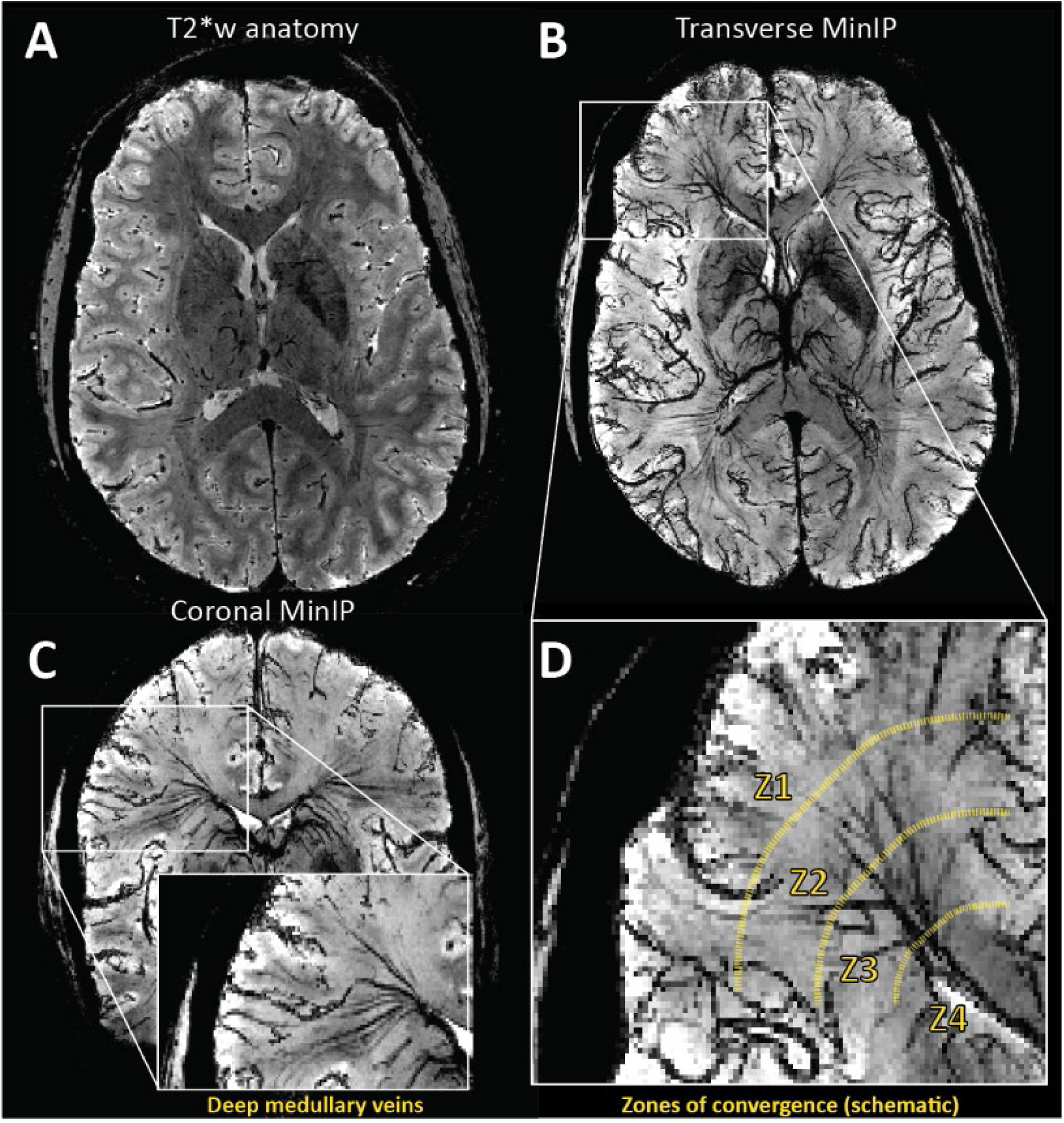
High-resolution T2*-weighted anatomical image of a representative subject. Based on this volume, a minimum intensity projection image (MinIP) is generated using 9.5mm slabs. The transverse and coronal MinIP images are shown in figure 1B and 1C, respectively. The inlay in 1C highlights the radial pattern of the large medullary veins. These veins fan outward from the ventricles to penetrate deeper white matter tissue. The expanded image in figure 1D highlights branching patterns that can be attributed to distinct convergence zones. As described by Okodura et al.^8^, zone 1 is characterized by the presence of superficial medullary veins and smaller branching patterns known as coat-rack and bamboo-branch unions. These structures are too fine to be resolved using MR imaging. The transition to zone 2 is mainly characterized by the candelabra structures created where laterally running tributaries converge with deeply running medullary veins. These deep medullary veins course through the third convergence zone to form palm-like unions with subependymal veins forming the fourth zone of convergence at the lateral ventricle.

The deep venous architecture has two implications concerning the WM BOLD-CVR contrast. The first being that large vessel density increases when moving inwards towards the ventricles, and the second being that aggregation of dHb through convergence in deep medullary veins will lead to BOLD-CVR signal delays and/or dispersion. While the grey matter BOLD-CVR response represents a robust surrogate for CVR mediated blood flow changes, the white matter response is nuanced and reflects multi-component behavior rooted in the initial control vessel response (i.e. true CVR) and then the downstream pooling behavior of the blood. If this is the case, then it obliges extra consideration when attributing changes in WM BOLD-CVR response shape, magnitude, and rate as possible pathological mechanisms.

To further investigate this notion, the vascular responses generated using hypercapnic-BOLD imaging were contextualized using high-resolution T2^*^ weighted imaging at 7 tesla. In addition to standard CVR maps generated using a controlled CO_2_ stimulus, maps of hemodynamic lag, lag-corrected CVR, signal onset, and dispersion were created. Parameter maps were compared with minimum intensity projections (MinIP) of venous structures derived from anatomical scans. This was driven by the hypothesis that differences in parametric response measures would spatially correlate with venous features described above. Finally, maps were registered to MNI space and correlated against a high-resolution medullary vein frequency atlas^15^.

## METHODS

This was a retrospective study in which a survey of the local imaging database identified datasets containing high-resolution dynamic BOLD imaging data (with consistent acquisition parameters) during which a controlled hypercapnic stimulus was administered. Further inclusion criteria were that participants also underwent a high-resolution T2^*^ weighted anatomical scan such that comparisons could be made between hemodynamic parameter maps and deep white matter vasculature. A total of 8 datasets were identified (average age 31 yrs., range 19-48 yrs., 4 females). Data was acquired with approval by the medical research ethics committee of University Medical Center Utrecht and written informed consent was obtained from all subjects. The experiments were performed according to the guidelines and regulations of the WMO (Wet Medisch Wetenschappelijk Onderzoek) and conforming to the declaration of Helsinki. All datasets have been used, at least in part, in previous publications^16, 17^. In this work, a series of new and original (re-)analyses were performed.

### Data Acquisition

MRI acquisition was done using a Philips 7 tesla MRI scanner (Philips, Best, The Netherlands) using a 32 channel receive coil in combination with a volume transmit coil (Nova Medical, Wilmington, MA, USA). Third-order image-based shimming was performed. Imaging data consisted of a high-resolution 3D multi-shot GE-EPI T2^*^ weighted anatomical acquisition^18^ (flip angle: 24°, TR/TE: 77/27 ms, EPI factor: 13, SENSE factor RL/FH: 2.3/1, reconstructed resolution: 0.5 mm isotropic, FOV: 240 × 150 × 192 mm^3^, acquisition matrix: 480 × 381 × 300 mm^3^, scan duration: 385 s) and a dynamic Blood Oxygenation Level Dependent (BOLD) acquisition (Multi-slice single-shot GE-EPI BOLD images (flip angle: 90°, TR/TE 3000/25 ms, EPI/SENSE factor 47/3, reconstructed resolution: 1.5 × 1.5 mm^2^, slice thickness: 1.6 mm, FOV: 217.6 × 192 mm^2^, acquisition matrix: 133 × 120, slices: 43). Arterial gases were manipulated during the BOLD scan using a third-generation RespirAct system (Thornhill Medical, Toronto, Canada) in combination with a rebreathing facemask. Two hypercapnic respiratory protocols were implemented within the included datasets. In three datasets a single 120s hypercapnic stimulus was administered interleaved with 120 second baseline periods. The remaining datasets used a respiratory paradigm consisting of 90s baseline periods interleaved with 90s hypercapnic, hyperoxic and hypercapnic-hyperoxic blocks, respectively. While it has been shown that the nature of the vasoactive stimulus can modulate hemodynamic parameter maps, this work aimed to evaluate the spatial distribution of various responses. Considering this, the inclusion of data derived using different hypercapnic paradigms was considered acceptable.

### Data Processing

Initial processing of the BOLD data consisted of brain extraction (BET^19^), motion correction (MCFLIRT^20^), and segmentation (FAST^21^) using FSL (FMRIB, Oxford, UK)^22^. Motion corrected BOLD data was then ‘scrubbed’ using functions included in the seeVR toolbox (seeVR, Utrecht, The Netherlands)^23^. This data cleaning step (outlined in figure 2) involved using a general linear model (GLM) to remove nuisance signals identified using motion parameters and their derivatives (time derivative, square), a linear drift term (1^st^ order Legendre polynomial), and the individual PetO_2_ trace measured by the RespirAct. Regressing out the O_2_ information ensured that further analysis was weighted primarily towards the vasodilatory effect of the hypercapnic stimulus. The regression of the PetO_2_ information was not included for the three normoxic datasets in which only a hypercapnic stimulus was applied. The associated Pet CO_2_ trace was convolved with a series of six double-gamma HRF functions (see equation 1: α_1_ = 1, β_1_ = 1-6, α_2_ = 1, β_2_ = 1)^24^ with increasing dispersion properties to simulate the ‘spreading-out’ of the WM BOLD-CVR response to step-changes in arterial CO_2_^7^; these convolved traces were then included in the GLM as explanatory variables. After running the GLM, the sum of the variance explained by the nuisance regressors was subtracted from the input data to generate the ‘cleaned’ BOLD data to be used for further analysis.

**Figure 2A:**
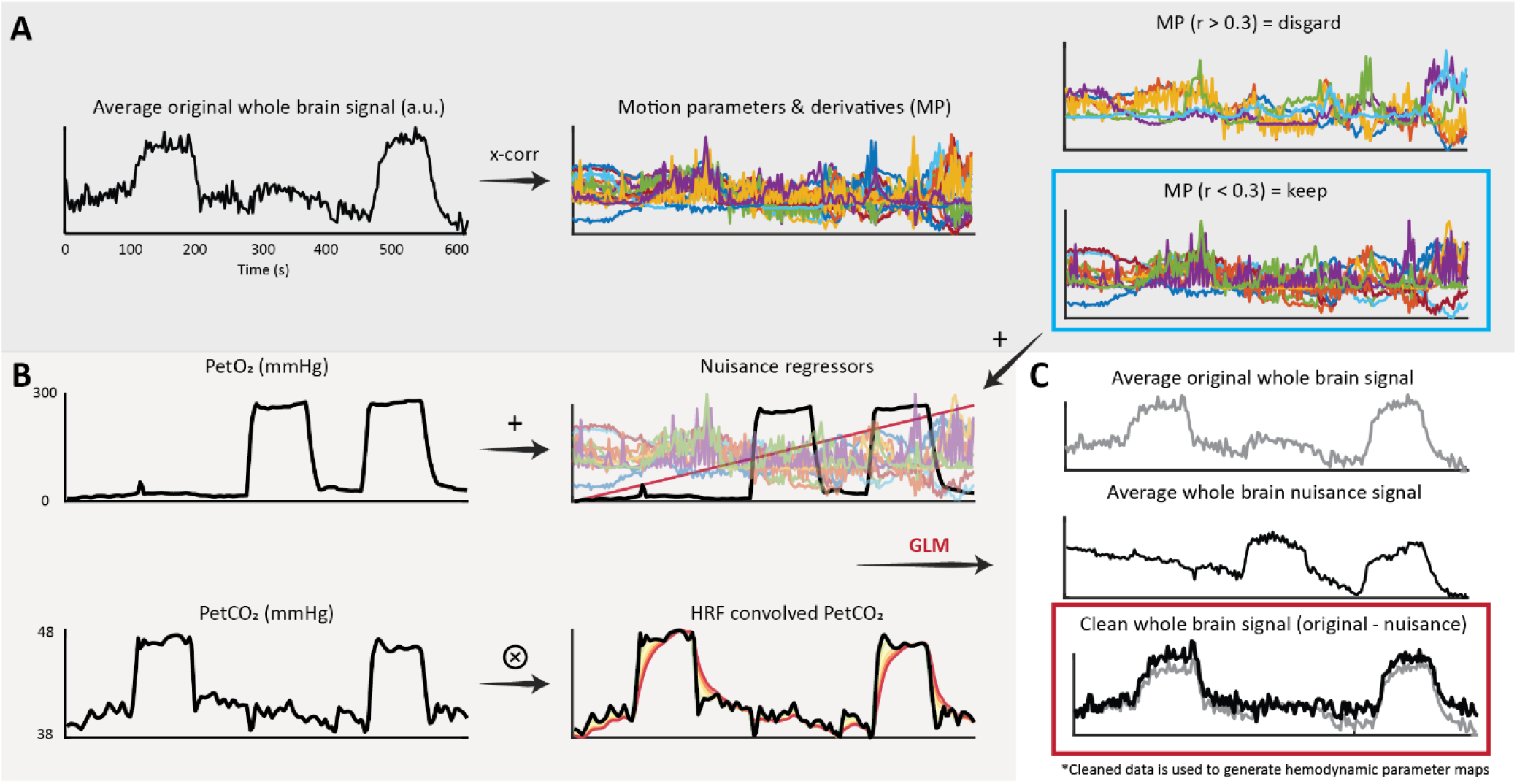
Occasionally motion parameters derived using intensity-based realignment methods can correlate with the hypercapnic task. This is in addition to subject motion related to hyperventilation or the transitions between baseline and hypercapnic breathing. In this case, highly correlated nuisance signals should not be included in the GLM since they will account for some of the desired signal responses. The average whole-brain signal is cross-correlated against the motion parameters and their derivatives. Parameters with an absolute correlation higher than 0.3 were rejected, while the remaining parameters are used for data scrubbing; Figure 2B: In 5 of 8 subjects, hyperoxic blocks were administered during the breathing protocol. In these datasets, the PetO_2_ traces were added as nuisance regressors to remove variance explained by the O_2_ challenge. A linear term was added to account for drift. The PetCO_2_ along with 6 HRF-convolved PetCO_2_ traces were included to provide a priori information regarding the vascular challenge; Figure 2C: nuisance regressors and data probes were used to explain the voxel-wise BOLD responses. For each voxel, the sum of the variance explained by the nuisance regressors was removed from the original voxel signal. The resulting ‘cleaned’ data was then used for further hemodynamic analysis.

### Hemodynamic lag mapping

Hemodynamic lag maps were generated based on a modified RAPIDTIDE approach^25, 26^ implemented in the seeVR toolbox. First, a manual bulk alignment between the PetCO_2_ trace and the average GM BOLD signal was performed. This manual approach minimized alignment errors that can occur due to noise or spike artifacts when using automated cross-correlation. The BOLD time-series data and PetCO_2_ traces were then linearly interpolated by a factor of 4 (effective TR: 0.75 s). The PetCO_2_ trace was then used as a seed to generate an optimized BOLD signal regressor^26^. GM voxels within a time-shift of −1 to 2 TRs (−3 to 6 seconds) and showing a correlation of <0.7 were temporally aligned and principal component analysis (PCA) was applied to identify the components that explained at least 85% of the signal variance. These principle components were then extracted to form a new seed trace. This process was iterated until the root mean squared error (RMSE) between subsequent traces converged to less than 0.005. Next, the final trace (optimized BOLD regressor) was cross-correlated against each brain voxel and the temporal signal lag was determined based on the maximum correlation value.

### CVR mapping

To generate the standard base CVR map (i.e. without accounting for WM hemodynamic lag), the bulk-aligned PetCO_2_ trace was regressed against each voxel of the baseline-normalized cleaned BOLD data. The slope of this linear regression was taken as the CVR in units of percent signal change per mmHg increase in CO_2_. Corrected CVR maps were generated by shifting the bulk-aligned PetCO_2_ trace according to the voxel-wise hemodynamic lag before performing the linear regression. Finally, a CVR difference map, from now on referred to as the ΔCVR map, was calculated by subtracting the base CVR map from the lag-corrected CVR map. Detailed reports of GM CVR from this (or a subset) data have been previously reported^5, 16^

### Onset and dispersion mapping

The double gamma function used to produce onset and dispersion maps is expressed as:

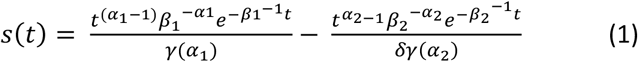

Where the α_1,2_ parameters model a global signal shift or onset (α_1_ is set equal to α_2_)^24^, β_1_ models the signal dispersion, β_2_ models the undershoot and δ modulates a vertical signal dispersion. For CO_2_ mediated BOLD responses, no undershoot is expected and so β_2_ was set equal to 1. To limit vertical dispersion, the δ parameter was set to 1000; this also restricted the overall contribution of the second gamma function for simplicity. The range of α_1,2_ was set from 1-2 in increments of 0.2 and the range of β_1_ was set from 1 to 200 in increments of 2.

Previously HRFs have been convolved with the PetCO_2_ trace to model the possible BOLD signal response^7^. However, it was apparent that the optimized BOLD signal regressor provided a more suitable archetype for the direct response of highly reactive vessels to changes in arterial CO_2_. Therefore, the optimized probe was taken as an initial reference point (instead of the PetCO_2_ trace) from which point the voxel-wise changes in onset or dispersion were investigated. In doing this, any modulations of the blood CO_2_ bolus (as defined by the PetCO_2_) occurring between the lungs and brain did not warrant consideration when modeling the brain response. This process generated a series of 600 HRF functions that were fit to each voxel time-series using a least-squares method. The onset and dispersion parameters correspond to the HRF with the highest R^2^ were then chosen to generate onset and dispersion maps. Example HRF functions and their convolutions are provided in supplementary figure 1.

### Image processing

Hemodynamic parameter maps were co-registered to the T2^*^ anatomical image via the associated mean BOLD image using a linear transformation (FLIRT: 9 parameters, mutual information, trilinear interpolation). Processing of the T2^*^ anatomical data consisted of brain extraction (BET) and segmentation with bias field removal (FAST). The bias-field removed T2^*^ image was then used to generate a minimum intensity projection (MinIP) image to highlight venous structures. For this, the minimum intensity value through a stack of 8 slices above and below each slice (17 slices, 9.5mm slab) of interest was projected. It should be noted that using the bias-field corrected image made little difference as compared to using the original T2^*^.

The T2^*^ anatomical image was then registered to a 1mm T2-weighted version of the MNI152 atlas^27^ using an affine registration with 12 degrees of freedom (FLIRT) followed by a non-linear registration (FNIRT^28^). Both the linear transformation matrix and non-linear warp fields were then applied to the T2^*^-registered hemodynamic parameter maps to bring them to MNI space. All maps were subsequently averaged such that the WM parameters could be evaluated in the context of medullar vein frequency using a high-resolution medullary vein atlas^15^. This atlas is expressed in units of ‘counts’ and is based on a dataset of 30 subjects scanned using the same 7 T scanner and a similar T2^*^-weighted sequence as the one used to acquire the data presented in this study.

### Statistical analysis

Considering the aim to investigate the spatial correlation between white matter venous topology and hemodynamic responses, the distributions of MNI-registered parameter maps were compared with the vessel frequency expressed in the medullary vein atlas. WM regions with significant medullary vein content were isolated in a 20mm slab beginning at the top of the ventricles (between slices 102-122 in the MNI volume; see supplementary figure 2). Heat-scatter plots of hemodynamic parameters versus vein frequency were generated and the Pearson correlation coefficient (rho, r) for overlapping voxels was determined. For correlation calculations involving the medullary atlas, a second atlas was generated in which all WM regions were included. This served as a proxy to evaluate relationships in voxels with presumably low vessel density (i.e. first to second zones of convergence; figure 1) that were not encompassed by the medullary mask. Correlation values below 0.39 were considered weak. Values from 0.4-0.59 were considered moderate. Values from 0.6 and 0.79 were considered strong and values above 0.8 were considered very strong. A supplementary correlation analysis was performed to examine the relationships between different hemodynamic parameters.

## RESULTS

The average baseline PetCO_2_ value across all participants was approximately 34.8 mmHg. The average PetCO_2_ value during the hypercapnic stimulus across all participants was 44.4 mmHg (for the five participants that experienced two hypercapnic blocks, the average value of both was taken). The average increase in PetCO_2_ between baseline and stimulus periods across all participants was approximately 9.5 mmHg.

Both hemodynamic lag analysis and signal dispersion modeling highlighted heterogeneous temporal responses in the WM in concordance with previous reports^5-7^. Visual inspection of single-subject data indicated higher lag and dispersion values localized centrally in regions above the ventricles and dispersing radially at points parallel to the ventricles (see figures 3 and 4). Moving towards the sub-cortical white matter, lag and dispersion values became lower. This was also the case for the calculated ΔCVR that generally associated spatially with the lag/dispersion. In the context of the venous topology, ΔCVR, lag, and dispersion were lower around the assumed first zone of convergence until the transition regions between the first and second zones. This is emphasized in figure 3A (white arrows), where both the base CVR and lag-corrected CVR remain low. When comparing these regions against the MinIP, there appear to be no large visible veins.

**Figure 3A:**
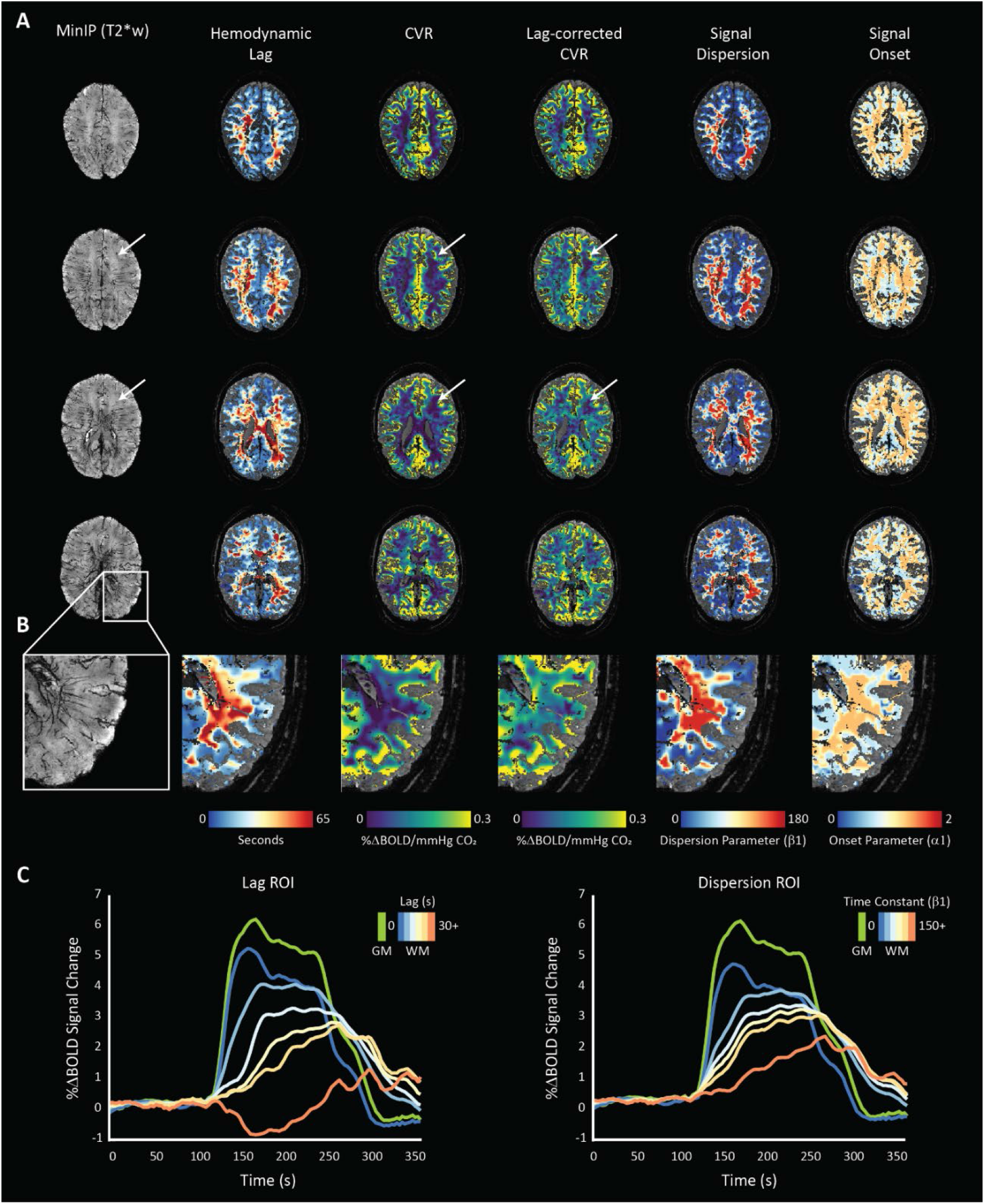
the MinIP and related hemodynamic parameter maps for a representative subject (48yrs, female) undergoing the normoxic, single hypercapnic-block paradigm are shown. Increases in lag-corrected CVR spatially correlate with longer hemodynamic lag and higher signal dispersion. Moreover, a clear relationship between lag and dispersion can be seen, as expected. White arrows indicate low CVR regions that remain unaffected when correcting for signal lag. When comparing to the corresponding region in the MinIP, this area seems devoid of large vessels suggesting low BOLD CNR related to low blood volume; Figure 3B: an enlarged region of the left-posterior WM where spatial correlations between medullary vessels (left: MinIP) and various hemodynamic parameters can be inferred; Figure 3C: the normalized BOLD time-series calculated within several regions of interest defined by increasing lag (left) and dispersion (right) are shown with a GM time-series for reference (green). These time courses highlight the relationship between increasing temporal parameters and the shape of the BOLD-CVR response.

**Figure 4:**
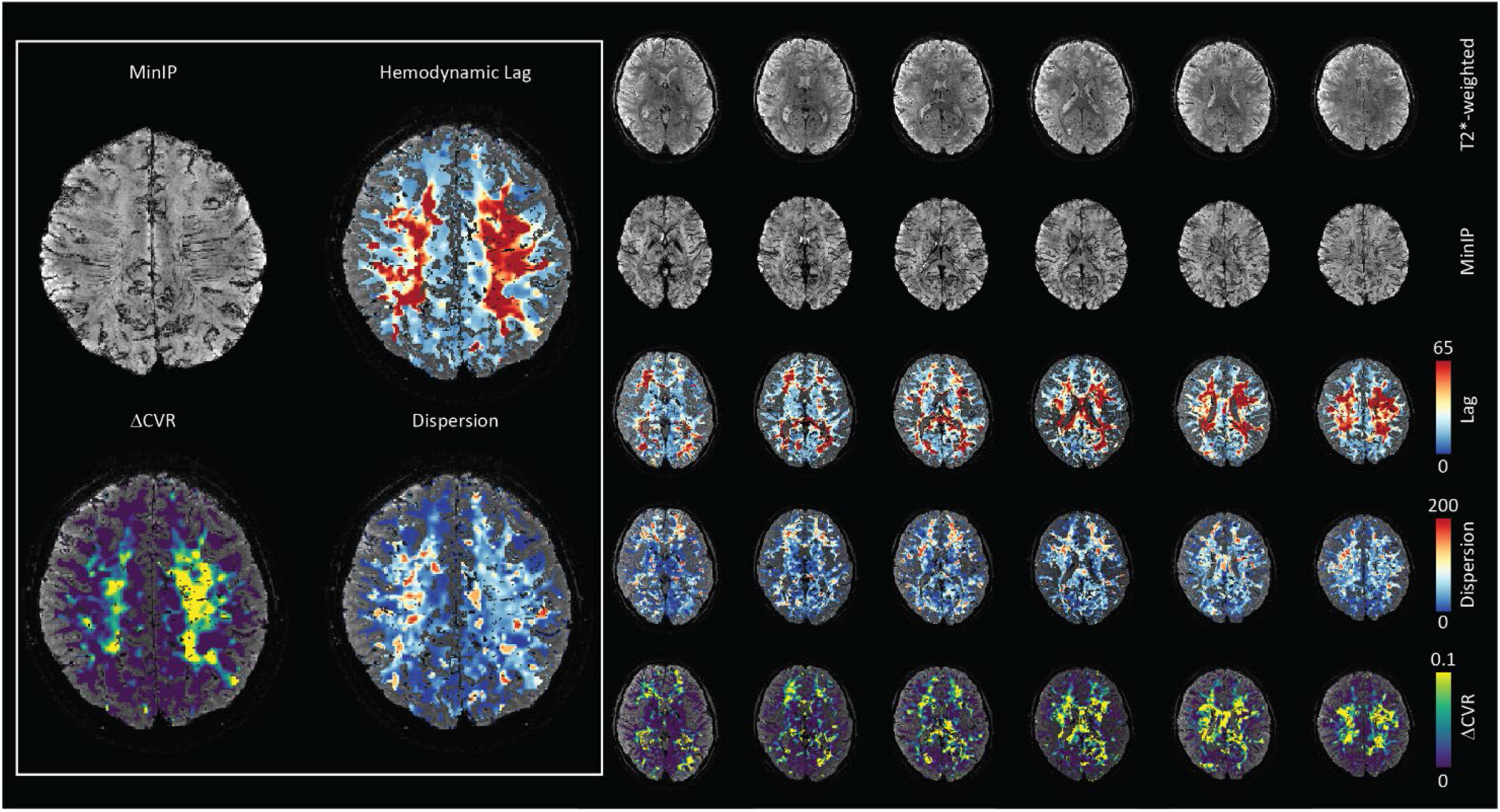
the MinIP and related hemodynamic parameter maps for a representative subject (19yrs, female) showing white matter lag, dispersion, and CVR difference maps. Spatial correlation between regions of temporally dispersed signals coincide with a larger magnitude in the corrected CVR. These are also regions with higher medullary vessel density as can be seen in the left inset.

Spatial patterns observed in single subject datasets remained consistent in the MNI-averaged maps (figure 5). The Pearson’s correlation value calculated between ΔCVR and the medullar frequency was 0.52 (0.58 when including all WM voxels). When comparing hemodynamic lag, the correlation coefficient was 0.42 (0.54 when including all WM voxels). For the dispersion and onset parameters, the Pearson correlation values were 0.44 (0.53 when including all WM voxels) and 0.43 (0.52 when including all WM voxels), respectively. Therefore, all hemodynamic parameters considered showed a moderate correlation indicating positive relationships with the frequency of medullary veins (figure 5).

**Figure 5A:**
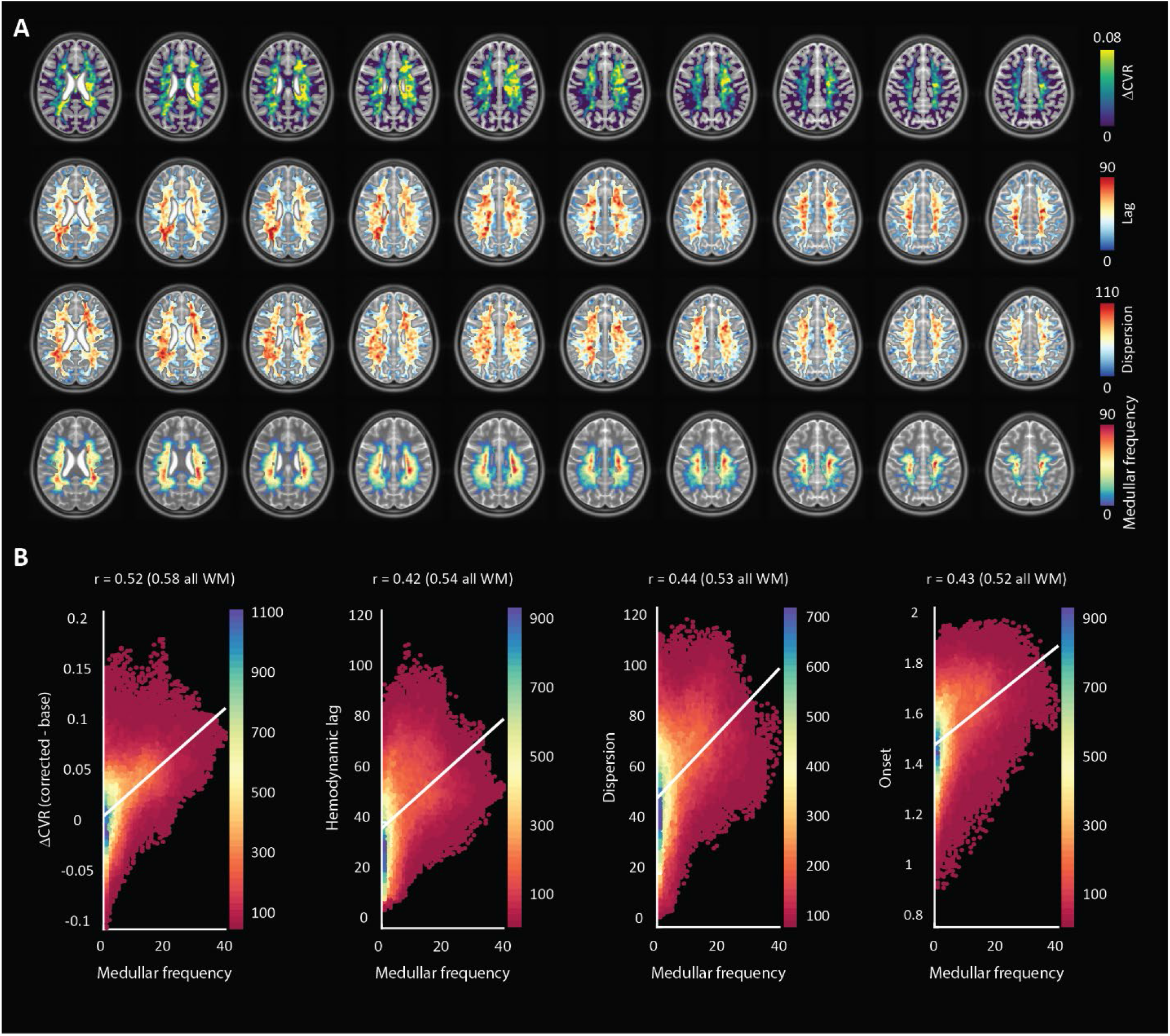
MNI-averaged (1 mm) hemodynamic parameter maps for all subjects showing ΔCVR, hemodynamic lag, and dispersion as well as the MNI-averaged medullar vein frequency atlas (bottom). Hemodynamic parameter maps were smoothed using a Gaussian kernel with a filter width of 5 voxels and an FWHM of 3mm; Figure 5B: heat-scatter plots showing the correlation between each parameter map with the medullar frequency map. A bin size of 50 was used and the color map represents the data counts for each bin. A depiction of the MNI slab included for correlation analysis along with heat-scatter plots comparing various hemodynamic parameters with one another are supplied in supplementary figure 2.

A secondary analysis was performed to compare the relationships between various hemodynamic parameters within the region defined by the medullary atlas. The Pearson value when comparing the dispersion and ΔCVR parameters against the hemodynamic lag showed a strong correlation at 0.70 and 0.75, respectively. Finally, the lag-corrected CVR showed a very strong correlation with the base CVR with a Pearson value of 0.94. This was expected, but highlights changes that can occur to CVR after correcting for lag. Scatter plots are provided in supplementary figure 2B.

A two-tailed t-test performed for each correlation showed that all were significantly different from the null hypothesis of zero correlation (p-value > 0.01).

## DISCUSSION

In this retrospective work, high-resolution T2^*^-weighted anatomical and functional imaging was applied to investigate the properties of the WM BOLD-CVR response to a vasoactive stimulus. The application of advanced hemodynamic analysis provided parametric maps of voxel-wise CVR, hemodynamic lag, lag-corrected CVR, signal onset and dispersion, and finally the ΔCVR. The main finding reported herein was that parametric maps showed moderate positive correlations with the frequency of the larger medullary veins responsible for draining much of the WM tissue. In light of the physical principles underpinning the BOLD signal contrast, this finding supports the hypothesis that drainage topology plays an important role in determining WM BOLD-CVR characteristics that might otherwise be attributed solely to auto-regulatory dilation or constriction.

For healthy subjects, CBF^29^ and CMRO_2_^30, 31^ are considerably lower in WM as compared to GM. Several studies have reported non-significant differences in the oxygen extraction fraction (OEF) between WM and GM in healthy subjects (see meta-analysis presented in Fan et al. 2020 ^32^) indicating a regional equilibrium between CBF and CMRO_2_. It follows that changes in venous hemoglobin saturation, along with venous CBV, will drive the WM BOLD signal contrast under hypercapnia. Moreover, as shown with ASL-based methods^29^, relatively low perfusion means that the WM BOLD contrast-to-noise (CNR; or detection sensitivity) is limited compared to GM, even at high magnetic field strength. While factors related to the true arterial response to CO_2_ (i.e. speed, flow distribution, CO_2_ sensitivity) will determine the venous hemodynamic input conditions, the accumulation of dHb in progressively larger collecting veins will modulate the WM BOLD-CVR response shape. A corollary is the impact of large draining veins that can reduce the specificity of cortical fMRI responses. Supplementary figure 3 provides an example of signal dispersion seen when comparing the total GM BOLD response to the response measured in a downstream region of the superior sagittal sinus. In the case of certain WM regions, large draining veins rather than control vessels may dominate response properties leading to ag and dispersion. This is partly due to due to the overall low vascular density in WM tissue. A conceptual explanation of how WM drainage properties influence the CVR response is shown in figure 6.

**Figure 6A:**
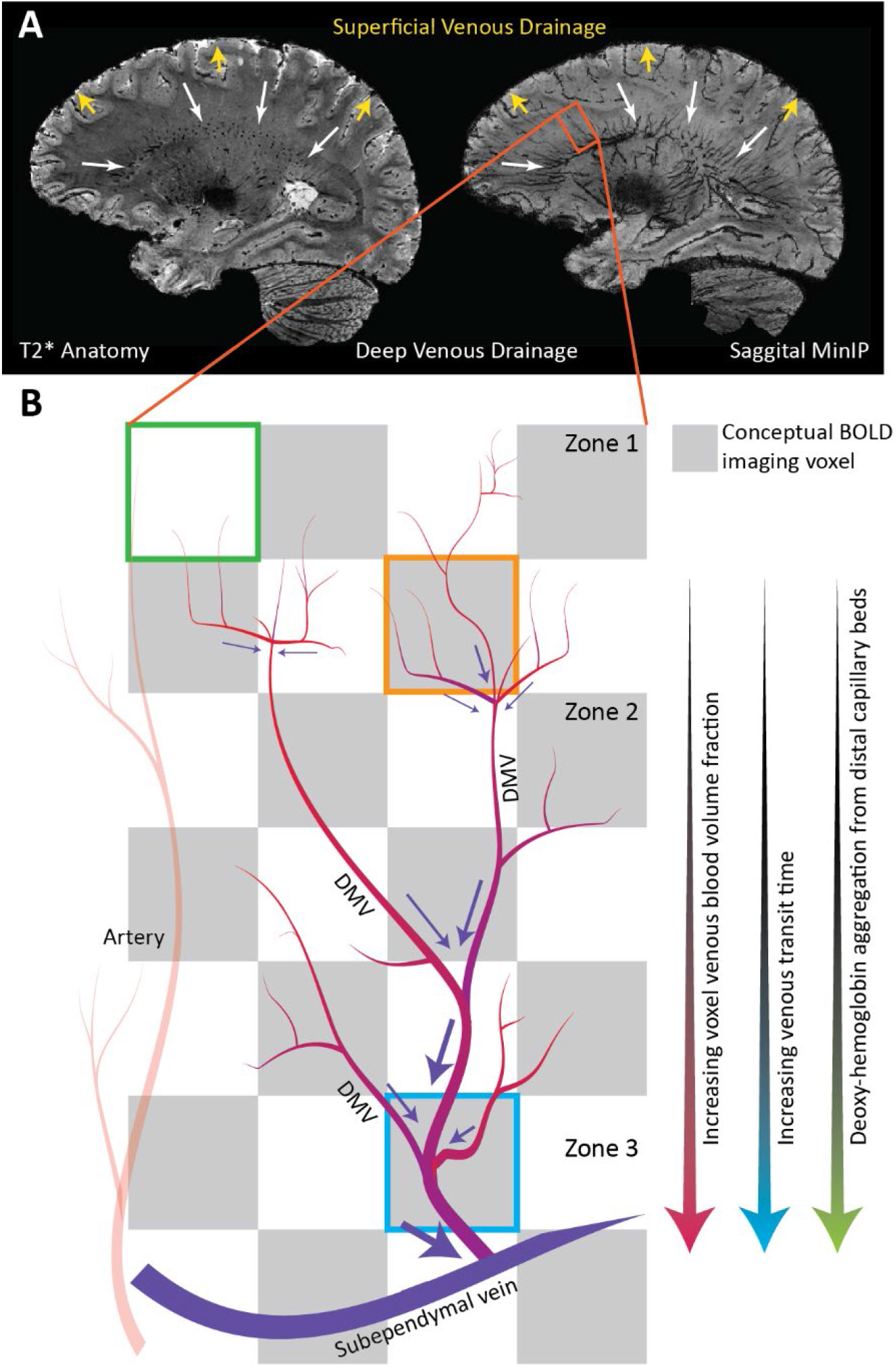
A sagittal slice from a representative subject T2*-weighted image. The corresponding MinIP is shown on the right. The venous drainage system can be separated into superficial and deep drainage networks. The superficial network drains cortical blood towards the superior sagittal sinus while the deep system drains white matter blood via the subependymal veins. The degree of collateral drainage between superficial and deep venous networks remains generally unclear. Figure 6B: The deep drainage architecture may have implications on the WM BOLD-CVR response to hypercapnia. In this conceptual representation, three representative BOLD voxels are considered. The green voxel contains very little venous blood volume. Here, reactive WM capillary beds may not evoke a strong BOLD signal response mainly due to the low total dHb content. Blood from such regions at the WM periphery join via tributaries at the second convergence zone (orange voxel). Here, the venous blood volume fraction increases appreciably, setting up the possibility to evoke stronger BOLD signal contrast. A similar effect occurs around zone 3 where visibly large medullary vessels come together. Moving inwards towards the ventricles, the total venous volume and total dHb content increase considerably. The venous transit time along with the congregation of blood from distal regions will lead to signal delays and dispersion. These effects are underscored when looking at signal increases in CVR maps after correcting for hemodynamic lag (figure 5).

Progressive pooling of dHb as the blood drains through the venous tributaries towards the subependymal veins has two important consequences. First, the total amount of dHb increases as blood moves away from superficial WM (below the sub-cortical WM) towards the larger peri-ventricular medullary veins. In these central regions, both the blood volume fraction and total dHb concentration may evoke a stronger BOLD signal effect as increased CBF reduces the OEF at capillary beds. Second, as the venous blood drains, it moves away from the actual site of arterial reactivity leading to a potential delocalization of the CVR signal response. Particularly since intravenous signal contributions from large veins medullary can dominate the BOLD signal response at lower fields^33^. This effect can be exacerbated by partial volume effects due to large voxels sizes (∼3 mm^3^) and large smoothing kernels (5-8mm^2^) that are typically applied in CVR studies using clinical 3T MRI systems.

In light of this new perspective, integrating knowledge of venous organization may aid in sharpening the application of CVR as a biomarker for diseases affecting the WM. Particularly in the case of medullary veins, whose characteristics have been shown to provide prognostic information in patients that have suffered from stroke^34^, and have been linked to a variety of disorders^10^. It is important to distinguish the absence of a BOLD-CVR response due to low perfusion or low blood volume from arterial aspects or actual impairments in the vascular ability to respond to a stimulus. Techniques to achieve might include scans that are sensitive to blood volume, or the rate-of-change in blood volume. For example, displacement encoding via stimulated echoes (DENSE)^35, 36^ MRI might reveal whether fast volume changes occur long before the presumably dispersed (or lagged) BOLD effects. To better understand possible large vein contributions, the acquisition of complimentary high-resolution anatomical T2^*^-images or susceptibility-weighted images SWI is recommended.

### Limitations

The use of the HRF-convolved data probes (figure 2B) can aid in restricting the variance explained by nuisance signals. This analysis step was inspired by the approach of convolving a design matrix with an assumed HRF function when modeling stimulus-response in task-based fMRI. However, increasing the complexity of the GLM will influence the residual data used to generate hemodynamic maps (figure 2C). The same may hold for ambiguities related to the linear model used to account for possible drift in the MR signal and BOLD signal drift arising due to endogenous CO_2_ accumulation or increased minute ventilation. Correct nuisance signal regression is a subject of debate in the field of fMRI and remains an open question in the context of BOLD-CVR response to hypercapnia. One way to minimize potential errors in BOLD-CVR experiments is the inclusion of multiple hypercapnic periods in a run as presented by Poublanc et. al^7^ and as applied in five of the subjects in this study. Including multiple transitions can provide a means through which to ‘regularize’ the GLM fit and avoid mischaracterizing low CNR signals for excessively dispersed signals. Accordingly, the inclusion of two different CO_2_ breathing protocols in this study (one block versus two) may have increased the variance in the presented data.

Another source of variance is related to the use of the medullary atlas. While this atlas was produced using the same MR system and a very similar acquisition sequence, it was not specifically generated using the datasets included in this study, nor was it created with the property to distinguish smaller vessels located around the first and second zones of convergence. Considering that the focus of this work was on the spatial relationship between hemodynamic patterns and venous topology, and since the amount of available data in this retrospective study was limited, this variation was accepted.

Finally, as with cortical fMRI, acquisitions with higher spatial resolutions can reduce localization errors^37^ related to the WM BOLD-CVR response that may be dominated by larger medullary draining veins. Ultra-high field MR systems are appealing for this reason. For clinical MR systems, the loss in SNR can be mitigated by using longer TR and increasing the duration of hypercapnic periods for more signal averaging. This approach will come at the expense of temporal fidelity, which may be recovered by using high acceleration factors (again, at the expense of SNR).

## Conclusion

The confluence of arterial hemodynamics and BOLD signal characteristics weighted by venous architecture unrelated to smooth-muscle mediated dilation/contraction play a significant role in defining the WM BOLD-signal response to hypercapnia. This caveat should be taken into account when attributing diseases mechanisms and/or progression to presumed impaired WM BOLD-CVR.

## Acknowledgments

The author thanks Hans Hoogduin and Marielle Philippens for their mentorship and supervision during the period in which the data used in this study was acquired. Also thanks to Wouter Schellekens, Mario Gilberto Báez Yáñez and Jaco J.M. Zwanenburg for valuable discussions, and Hugo Kuijf for providing the medullar vein atlas.

## Data and code availability

The analysis tools used to generate the results presented in this manuscript are freely available via the open-source seeVR toolbox (https://github.com/abhogal-lab/seeVR). MRI and physiological data can be made available based on the submission of a formal project outline.

## References

1. Fisher JA, Sobczyk O, Crawley A, et al. Assessing cerebrovascular reactivity by the pattern of response to progressive hypercapnia. Human Brain Mapping. 2017; 38: 3415–27.

2. Sam K, Peltenburg B, Conklin J, et al. Cerebrovascular reactivity and white matter integrity. Neurology. 2016; 87: 2333–9.

3. Sam K, Crawley AP, Conklin J, et al. Development of White Matter Hyperintensity Is Preceded by Reduced Cerebrovascular Reactivity. Annals of Neurology. 2016; 80: 277–85.

4. Liem MK, Lesnik Oberstein SA, Haan J, et al. Cerebrovascular reactivity is a main determinant of white matter hyperintensity progression in CADASIL. AJNR American journal of neuroradiology. 2009; 30: 1244–7.

5. Bhogal AA, Philippens ME, Siero JC, et al. Examining the regional and cerebral depth-dependent BOLD cerebrovascular reactivity response at 7T. Neuroimage. 2015; 114: 239–48.

6. Thomas BP, Liu P, Park DC, van Osch MJP and Lu H. Cerebrovascular Reactivity in the Brain White Matter: Magnitude, Temporal Characteristics, and Age Effects. Journal of Cerebral Blood Flow & Metabolism. 2013; 34: 242–7.

7. Poublanc J, Crawley AP, Sobczyk O, et al. Measuring cerebrovascular reactivity: the dynamic response to a step hypercapnic stimulus. Journal of cerebral blood flow and metabolism : official journal of the International Society of Cerebral Blood Flow and Metabolism. 2015; 35: 1746–56.

8. Okudera T, Huang YP, Fukusumi A, Nakamura Y, Hatazawa J and Uemura K. Micro-angiographical studies of the medullary venous system of the cerebral hemisphere. Neuropathology : official journal of the Japanese Society of Neuropathology. 1999; 19: 93–111.

9. Khalatbari H, Wright JN, Ishak GE, Perez FA, Amlie-Lefond CM and Shaw DWW. Deep medullary vein engorgement and superficial medullary vein engorgement: two patterns of perinatal venous stroke. Pediatric radiology. 2021; 51: 675–85.

10. Taoka T, Fukusumi A, Miyasaka T, et al. Structure of the Medullary Veins of the Cerebral Hemisphere and Related Disorders. Radiographics : a review publication of the Radiological Society of North America, Inc. 2017; 37: 281–97.

11. Huang YP, Okudera T, Fukusumi A, et al. Venous architecture of cerebral hemispheric white matter and comments on pathogenesis of medullary venous and other cerebral vascular malformations. The Mount Sinai journal of medicine, New York. 1997; 64: 197–206.

12. Duvernoy HM, Delon S and Vannson JL. Cortical blood vessels of the human brain. Brain research bulletin. 1981; 7: 519–79.

13. Vigneau-Roy N, Bernier M, Descoteaux M and Whittingstall K. Regional variations in vascular density correlate with resting-state and task-evoked blood oxygen level-dependent signal amplitude. Human Brain Mapping. 2014; 35: 1906–20.

14. Bernier M, Cunnane SC and Whittingstall K. The morphology of the human cerebrovascular system. Human Brain Mapping. 2018; 39: 4962–75.

15. Kuijf HJ, Bouvy WH, Zwanenburg JJ, et al. Quantification of deep medullary veins at 7 T brain MRI. European radiology. 2016; 26: 3412–8.

16. Bhogal AA, Siero JC, Fisher JA, et al. Investigating the non-linearity of the BOLD cerebrovascular reactivity response to targeted hypo/hypercapnia at 7T. Neuroimage. 2014; 98: 296–305.

17. Champagne AA and Bhogal AA. Insights Into Cerebral Tissue-Specific Response to Respiratory Challenges at 7T: Evidence for Combined Blood Flow and CO(2)-Mediated Effects. Frontiers in physiology. 2021; 12: 601369.

18. Zwanenburg JJ, Versluis MJ, Luijten PR and Petridou N. Fast high resolution whole brain T2* weighted imaging using echo planar imaging at 7T. Neuroimage. 2011; 56: 1902–7.

19. Smith SM. Fast robust automated brain extraction. Hum Brain Mapp. 2002; 17: 143–55.

20. Jenkinson M, Bannister P, Brady M and Smith S. Improved optimization for the robust and accurate linear registration and motion correction of brain images. Neuroimage. 2002; 17: 825–41.

21. Zhang Y, Brady M and Smith S. Segmentation of brain MR images through a hidden Markov random field model and the expectation-maximization algorithm. IEEE transactions on medical imaging. 2001; 20: 45–57.

22. Smith SM, Jenkinson M, Woolrich MW, et al. Advances in functional and structural MR image analysis and implementation as FSL.Neuroimage. 2004; 23 Suppl 1: S208–19.

23. Bhogal AA. seeVR: a toolbox for analyzing cerebrovascular reactivity data. Zenodo. 2021; (v1.01).

24. Yao JF, Yang HS, Wang JH, et al. A novel method of quantifying hemodynamic delays to improve hemodynamic response, and CVR estimates in CO2 challenge fMRI. Journal of cerebral blood flow and metabolism : official journal of the International Society of Cerebral Blood Flow and Metabolism. 2021; 41: 1886–98.

25. Champagne AA, Bhogal AA, Coverdale NS, Mark CI and Cook DJ. A novel perspective to calibrate temporal delays in cerebrovascular reactivity using hypercapnic and hyperoxic respiratory challenges. Neuroimage. 2017.

26. Donahue MJ, Strother MK, Lindsey KP, Hocke LM, Tong Y and Frederick BD. Time delay processing of hypercapnic fMRI allows quantitative parameterization of cerebrovascular reactivity and blood flow delays. Journal of cerebral blood flow and metabolism: official journal of the International Society of Cerebral Blood Flow and Metabolism. 2016; 36: 1767–79.

27. Fonov VS, Evans AC, McKinstry RC, Almli CR and Collins DL. Unbiased nonlinear average age-appropriate brain templates from birth to adulthood. Neuroimage. 2009; 47: S102.

28. Jenkinson M, Beckmann CF, Behrens TE, Woolrich MW and Smith SM. FSL. Neuroimage. 2012; 62: 782–90.

29. van Osch MJ, Teeuwisse WM, van Walderveen MA, Hendrikse J, Kies DA and van Buchem MA. Can arterial spin labeling detect white matter perfusion signal? Magnetic resonance in medicine. 2009; 62: 165–73.

30. Paech D, Nagel AM, Schultheiss MN, et al. Quantitative Dynamic Oxygen 17 MRI at 7.0 T for the Cerebral Oxygen Metabolism in Glioma. Radiology. 2020; 295: 181–9.

31. Pantano P, Baron JC, Lebrun-Grandié P, Duquesnoy N, Bousser MG and Comar D. Regional cerebral blood flow and oxygen consumption in human aging. Stroke. 1984; 15: 635–41.

32. Fan AP, An H, Moradi F, et al. Quantification of brain oxygen extraction and metabolism with [15O]-gas PET: A technical review in the era of PET/MRI. Neuroimage. 2020; 220: 117136.

33. Kim S-G and Ogawa S. Biophysical and physiological origins of blood oxygenation level-dependent fMRI signals. Journal of cerebral blood flow and metabolism : official journal of the International Society of Cerebral Blood Flow and Metabolism. 2012; 32: 1188–206.

34. Yu X, Yuan L, Jackson A, et al. Prominence of Medullary Veins on Susceptibility-Weighted Images Provides Prognostic Information in Patients with Subacute Stroke. American Journal of Neuroradiology. 2016; 37: 423–9.

35. Adams AL, Kuijf HJ, Viergever MA, Luijten PR and Zwanenburg JJM. Quantifying cardiac-induced brain tissue expansion using DENSE. NMR in biomedicine. 2019; 32: e4050.

36. Sloots JJ, Biessels GJ and Zwanenburg JJM. Cardiac and respiration-induced brain deformations in humans quantified with high-field MRI.Neuroimage. 2020; 210: 116581.

37. Olman CA, Inati S and Heeger DJ. The effect of large veins on spatial localization with GE BOLD at 3 T: Displacement, not blurring. Neuroimage. 2007; 34: 1126–35.

