## supplementary figure for "Medullary vein architecture modulates the white matter BOLD cerebrovascular reactivity signal response to CO_2_: observations from high-resolution T2^*^ weighted imaging at 7T"

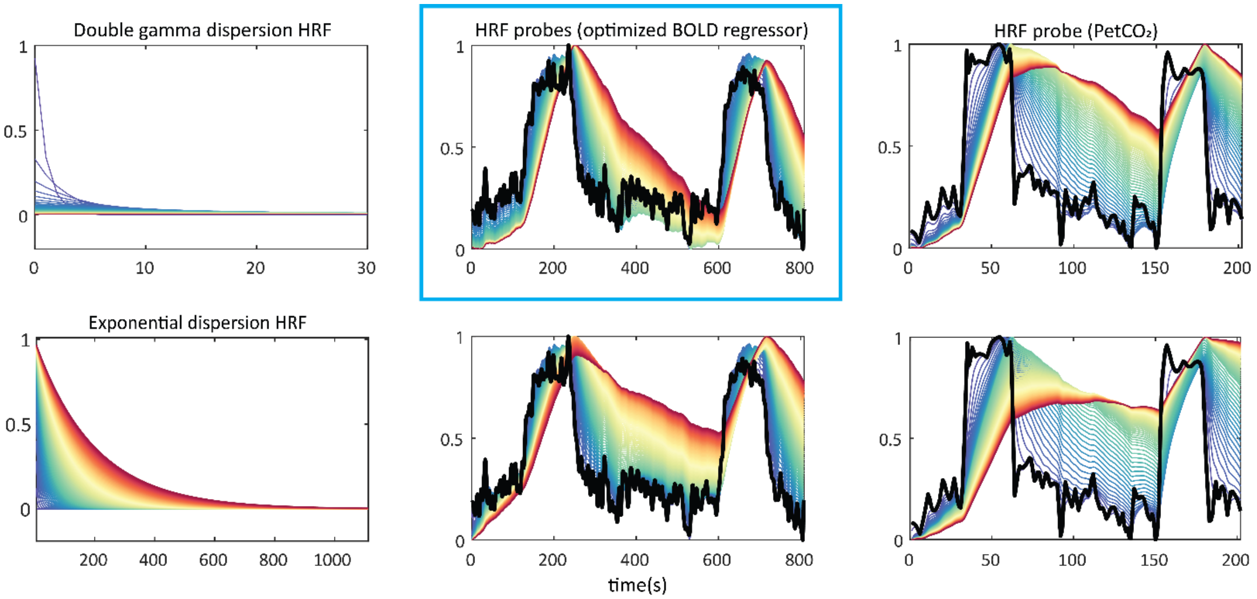


Supplemental Figure 1: In the left column, two different hemodynamic response functions with increasing dispersion time constants are shown. On top is the double-gamma function (Yao et al., 2011), and below is an exponential function (Poublanc et al., 2015). In the center column, each HFR is convolved with an optimized BOLD signal regressor that was generated while calculating the hemodynamic lag parameter map. This optimized trace represents the response generated by the most reactive vessels (or voxels) in the GM. As such, it may be considered as an indicator of how the brain 'senses' the incoming $CO_2$ bolus and thus, manifests the true physiological HRF. In the right column, each HRF has been convolved with the $PetCO_2$ trace (black). This approach assumes that one of the applied HRFs will provide a suitable model for both the vascular response as well any assumed physiological processes that occur between the lungs ($PetCO_2$) and the brain (BOLD response). For the dispersion analysis, the double gamma function convolved with the optimized BOLD regressor (blue box) was chosen since it utilized the most direct reference signal and allowed simultaneous modeling of the onset parameter, which can also potentially inform on $CO_2$ arrival.

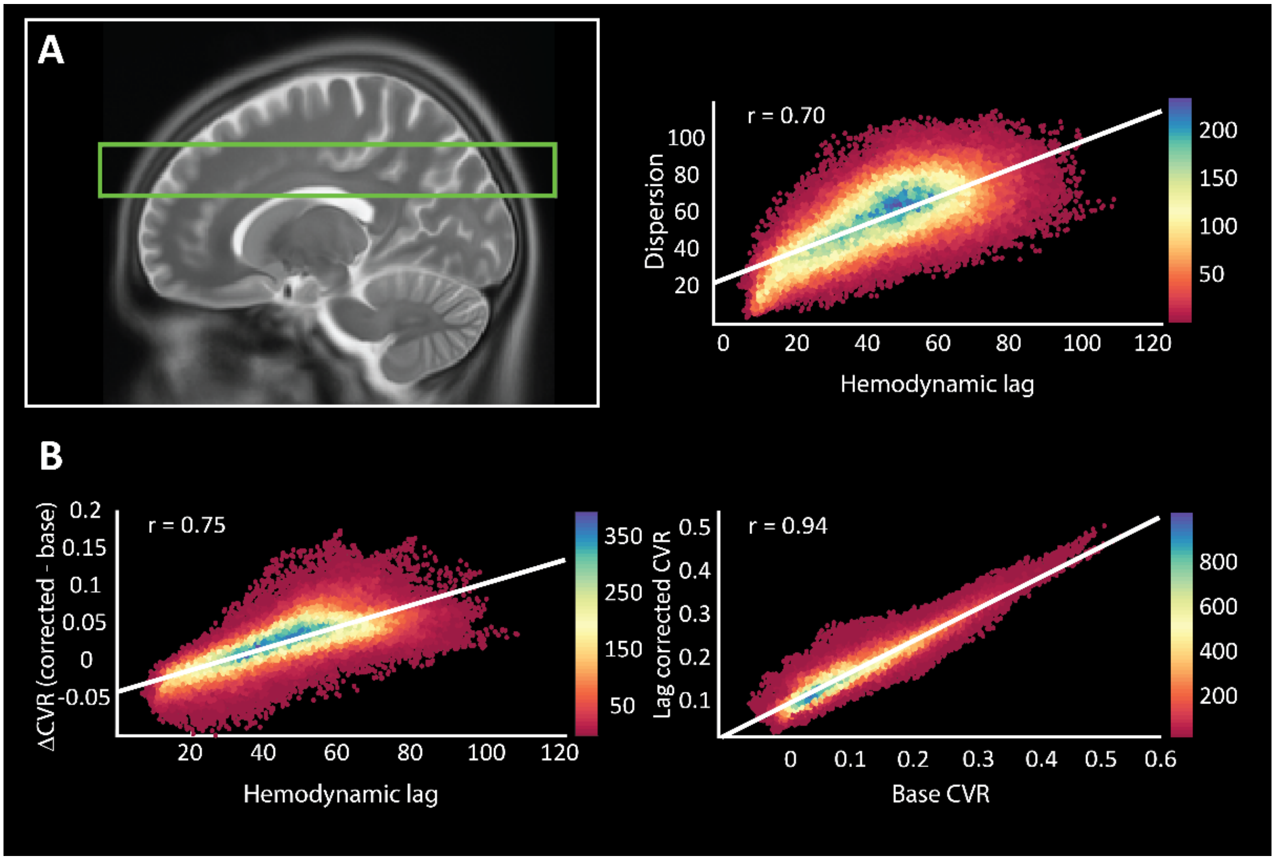


Supplemental figure 2A: The green box denotes the region of interest used when correlating the various hemodynamic parameter maps and the medullary vein atlas. Slices at lower locations contained contributions from the basal ganglia and insula, while slices at locations higher did not contain sufficient medullary vessel content; Figure 2B: heat-scatter plots show the relationship between various hemodynamic parameters. Dispersion and lag analysis entails two alternate methods for probing the same physiologal property. Namely, the delayed or slowed response to the abrupt hypercapnic stimulus. Therefore it is of no surprise that the correlation between these two parameters is strong (r = 0.7). The same can be said for base CVR and lag-corrected CVR (r = 0.94). Interestingly, a strong correlation was found between lag-corrected CVR and hemodynamic lag (r = 0.75) where the highest voxels counts are likely to represent regions around larger venous structures.

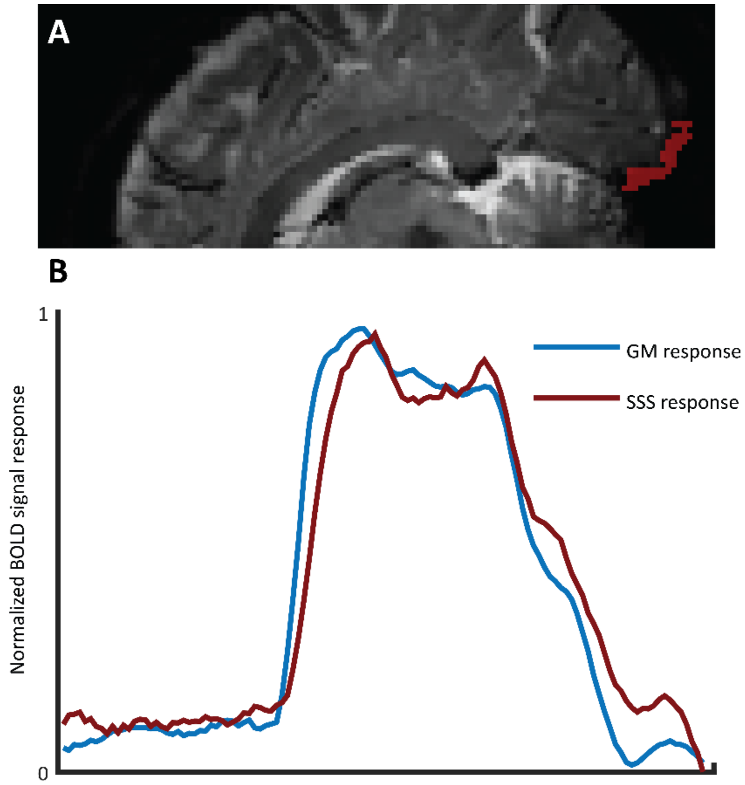


Supplemental figure 3A: A manually drawn mask containing voxels of the superior sagittal sinus (SSS) at a distal point to the rear of the cerebellum is shown. Figure 3B: The normalized average grey matter signal trace (blue) is compared with the normalized signal response contained with the SSS mask. A clear dispersion effect can be seen which will also manifest as an increased hemodynamic delay when performing lag analysis. This example serves as a validation for the notion that draining topology, and not direct vascular control, plays an important role in modulating certain BOLD-CVR responses. The data shown are from the same subject represented in figure 3 of the main manuscript.
